# A novel approach of human geroprotector discovery by targeting the converging subnetworks of aging and age-related diseases

**DOI:** 10.1101/326264

**Authors:** Jialiang Yang, Bin Zhang, Sander Houten, Eric Schadt, Jun Zhu, Yousin Suh, Zhidong Tu

## Abstract

A key goal of geroscience research is to discover effective interventions to extend human healthspan, the years of healthy life. Currently, majority of the geroprotectors are found by testing compounds in model organisms; whether these compounds will be effective in humans is largely unknown. Here we present a novel strategy called ANDRU (<u>a</u>ging <u>n</u>etwork based <u>dru</u>g discovery) to help the discovery of human geroprotectors. Instead of relying on model organisms, this approach is driven by human genomic and pharmacogenomic data. It first identifies human aging subnetworks that putatively function at the interface between aging and age-related diseases; it then screens for pharmacological or genetic interventions that may “reverse” the age-associated transcriptional changes seen in these subnetworks. We applied ANDRU to human adipose and artery tissues. In adipose tissue, *PTPN1*, a target for diabetes treatment and *APOE*, a known genetic factor for human longevity and diseases like Alzheimer’s disease, were ranked at the top. For small molecules, conjugated linoleic acid and metformin, a drug commonly used to treat type 2 diabetes, were ranked among the top compounds. In artery tissue, N-methyl-D-aspartate antagonists and curcumin were ranked at the top. In summary, ANDRU represents a promising human data-driven strategy that may speed up the discovery of interventions to extend human healthspan.

## Introduction

Aging is a major risk factor for age-related diseases (ARDs) and the ultimate cause for most human mortalities^1^. It has been demonstrated in model organisms, that genetic, environmental, and pharmacological interventions capable of extending lifespan are associated with delayed onset and progression of multiple age-related diseases^2^. Such observations have laid the foundations for the hypothesis of geroscience^1^, which states that aging is the major modifiable risk factor for most chronic diseases, and it is possible to simultaneously prevent multiple age-related diseases by targeting the basic biology of aging.

Through drug screening and testing in model organisms, more than four hundred anti-aging drugs have been identified^3,4^. Despite the progress, the development of human geroprotectors is facing significant challenges^5^. For example, since the majority of the geroprotectors were identified in model organisms with little human data support, their effectiveness in promoting human healthspan remains largely unknown. Such uncertainty poses high risk for the pharmaceutical industry to engage in the very costly clinical trials^5^.

Human aging is a complex process with a large number of genes and pathways being involved. For example, we and several other groups have shown that the expression of hundreds to thousands of genes can change with age in various tissues^6,7^. To achieve a holistic understanding of the aging process and to better elucidate the complex interconnections between aging and ARDs, systems and network approaches have been explored^8,9^. For example, Wang et al. showed that age-related disease genes were closer to aging genes in a protein-protein interaction (PPI) network^10^. We found that age-related disease categories shared functional terms including canonical aging pathways, suggesting that conserved pathways of aging might simultaneously influence multiple ARDs in humans^11^. We also developed a network algorithm called GeroNet and showed that it could prioritize biological processes mediating the connections between aging and ARDs^12^. Despite these progresses, most existing models neither considered tissue specificity nor incorporated the network dynamics associated with aging, and therefore, they captured human aging systems with rather limited accuracy. Although more recent work has started to address the tissue specificity for human aging research^13^, quantitative tissue specific aging network models are yet to be developed to allow more accurate characterization of the interplay between aging and ARDs in humans.

In addition to providing biological insights into aging and disease mechanisms, network biology can also facilitate drug discovery^14^. A promising area of development is based on the Connectivity Map (CMap) concept^15^, which uses gene expression responses to compound/gene perturbations for discovering the mode of action or drug repositioning^16^. An increasing number of studies have provided proof-of-the-concept and multiple successful applications have been reported, including a study in *C. elegans* to identify compounds that mimic the effect of caloric restriction to extend lifespan^17–19^.

Previously, we worked on a large human genomic dataset generated by the Gene-Tissue Expression (GTEx) project^20^. We showed that GTEx data allowed us to perform a comprehensive survey of tissue-specific aging mechanisms which recapitulated multiple hallmarks of aging^7^. In this study, we demonstrate a novel framework called ANDRU (<u>a</u>ging <u>n</u>etwork based <u>dru</u>g discovery) to further leverage this large human dataset to identify interventions that may help to extend human healthspan.

## Results

### An overview of the ANDRU pipeline

ANDRU takes tissue specific transcriptomic data from humans of young and old ages as input. It outputs a list of candidate compounds or target genes that may help to slow the aging and provide geroprotection in the corresponding tissues. A network model is applied to reveal the underlying subnetworks that may function as the interface between aging and ARDs, which will be used to prioritize target genes or compounds to achieve optimized geroprotection. An overview of ANDRU is shown in Fig 1. First, for each tissue, we construct gene co-expression networks^21^ from young and old tissue samples, respectively. The subnetworks (or modules) are constructed based on correlations among gene expressions and their topological overlaps. Since subnetworks often show enrichment for specific biological functions, it provides a data-driven approach of dividing the transcriptome into multiple smaller functional modules. Second, using aging differentially expressed genes (DEGs) and known disease gene sets, we prioritize subnetworks that are strongly influenced by both aging and diseases. Third, the aging DEGs in the prioritized subnetworks are used to query upon pharmacogenomic datasets to identify interventions that may “reverse” the age-associated expression changes in the corresponding subnetworks. The top ranked drugs or genes are further evaluated based on independent data and/or literatures.

**Fig 1.**
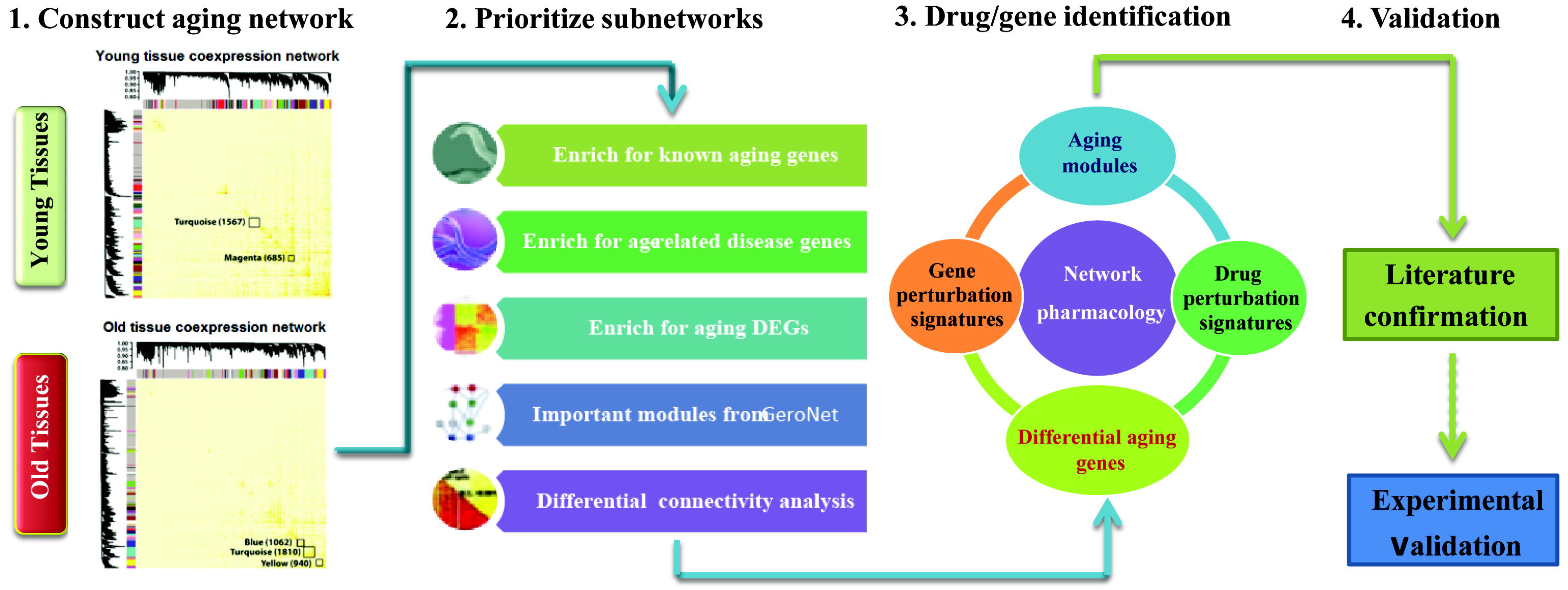
An overview of the ANDRU pipeline. Four major steps are 1). Tissue and age specific aging network construction; 2) Subnetwork prioritization based on multiple criteria to highlight the converging point of aging and age-related diseases; 3) Application of a pharmacogenomic approach to rank candidate drugs or genes. As an example, the down-regulated genes in drug 3’s perturbation signature is highly enriched for the up-regulated aging DEGs in the orange subnetwork, and therefore drug 3 is considered as a hit; and 4) Either literature or experimental based confirmation of the top candidate compounds or genes.

We first applied ANDRU to subcutaneous adipose tissue samples from GTEx. Adipose is a dynamic tissue showing profound changes related to aging and can play important roles in the development of multiple ARDs^22^. The GTEx data (v6p) profiled transcriptomes of 56,238 genes in 350 adipose tissue samples. According to donor’s chronological age, we classified 36 samples into the young group (age <= 35) and 52 samples into the old group (age >=65). The age and sex distributions of the samples used are shown in Fig S1.

### Deriving and comparing young and old adipose tissue gene co-expression networks

103 and 85 subnetworks are found in young and old adipose tissues respectively using WGCNA (weighted gene co-expression network analysis)^21^. The largest module in the old adipose is “turquoise” (denoted hereafter as turquoise^old^), which consists of 1,299 protein-coding genes (see Fig 2A). This module is significantly enriched for “mitochondrial matrix” (FDR =1.90E-13) and “oxidoreductase” (FDR=8.34E-13) (Table S1). Other large modules are enriched for “cell junction”, “Ubl conjugation pathway”, and “signal peptide” (see details in Table S2). The top modules in the young adipose tissue (see Fig 2B) include “turquoise” module (1,010 protein-coding genes), which is mostly enriched for “DNA repair”; “yellow” module (939 protein-coding genes), which is mostly enriched for “WD repeat”, and “magenta” module (604 protein-coding genes, denoted hereafter as magenta^young^), which is mostly enriched for “mitochondrion” (Table S2).

**Fig 2.**
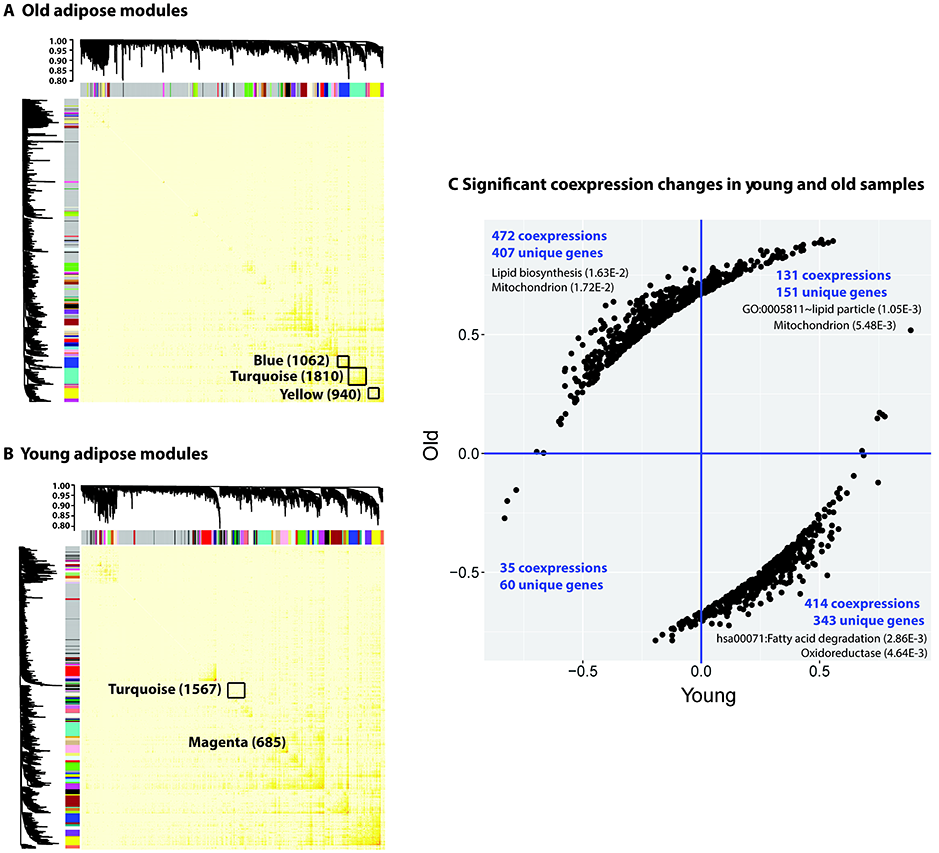
WGCNA networks in (A) young and (B) old adipose tissues, a few large size modules are highlighted with their number of genes annotated (both protein coding and non-protein coding genes are counted), and (C) the significant co-expression changes between young and old samples for the turquoise^old^ module.

Since WGCNA constructs networks from young and old samples separately, the subnetwork IDs (as color names) are assigned independently and do not match between the young and old networks. To map and compare the global network structures in young and old tissues, we considered gene overlap between the two networks. We found that turquoise^old^ most significantly overlapped with magenta^young^ with Jaccard index of 0.12 (see Table S3). Both modules are significantly enriched for mitochondria and transition peptide related functions. Since the magenta^young^ and turquoise^old^ show quite different sizes, this could imply that the co-expression wiring has changed substantially due to aging for genes in this subnetwork. To a further investigation, we performed a differential connectivity analysis^23^. Specifically, we aimed to identify the significant differentially co-expressed gene pairs between young and old networks. From this analysis, 1,052 significant differential co-expressions (p-value ≤ 5.0E-4) involving 643 unique genes were found (Fig 2C. and Dataset 1). The 643 genes are significantly enriched for “oxidoreductase” (FDR of 1.66E-6), “GO:0005759~mitochondrial matrix” (FDR of 5.26E-5), and “hsa00350:tyrosine metabolism” (FDR of 6.85E-3) (Table S4). For these significantly changed co-expression pairs, the majority of them show gained co-expression in the old network. For example, 708 gene-gene pairs showed significant correlations in old samples but no significant correlation in young samples. In contrast, only 38 gene-gene pairs that showed significant correlation in young samples are no longer strongly correlated in old tissues (loss of correlation). For the 505 genes that are involved in the 708 gene pairs that gained correlation in old adipose (regardless of the sign of the correlation), they are enriched for “mitochondrion” (FDR = 5.2E-5), “oxidoreductase” (FDR = 3.6E-5), and “fatty acid degradation” (FDR = 4.5E-2). For the 1,052 significant differential co-expressions, majority of them show changes in their co-expression directions. For example, there are 472 negative correlations in the young samples but become positively correlated in the old samples (in the 2^nd^ quadrant in Fig 2C). Similarly, 414 positive co-expressions in the young tissues became negatively correlated in the old tissues (in the 4^th^ quadrant in Fig 2C). This result indicates that substantial network re-wiring occurred in the adipose tissue during aging. The functions of the re-wired genes suggest potential link with age-dependent adipose function declines in insulin sensitivity, lipolytic, and fatty acid responsiveness^24^. Since we had imbalanced number of samples for young vs. old adipose tissues, to ensure the gain of co-expression was not due to larger sample size for old adipose, we performed a down-sampling calculation. Specifically, we randomly selected 36 samples from the old adipose tissue for 20 times, and calculated the co-expression values for the same 1,052 gene pairs. We found that the results were consistent with the ones obtained from using all the 52 old adipose samples (Dataset 1). Therefore, the gain of co-expression in the old adipose was not caused by the larger sample size.

### Prioritizing subnetworks that may function at the interface between aging and age-related diseases

As co-expression network divides transcriptome into a number of non-overlapping subnetworks, it is unclear if all these subnetworks are equal with respect to aging and ARDs. Based on our previous work^12^, we hypothesize that subnetworks are differentially involved in aging and make unequal contribution to the development of age-related diseases. To evaluate the relative importance of each subnetwork, we considered a five-criterion evaluation and applied it to the old tissue sample derived co-expression network. We chose old network for this evaluation since in general, ARDs mainly occur in older people; therefore, modules derived from the old samples are more likely to reveal the interconnection between aging and ARDs. We describe the details of each criterion below.

*Criterion 1. Enrichment for GenAge genes*: GenAge database (Build 18) contains 305 literature-based manually curated putative human aging genes^25^, and has been used as a reference aging gene list by several studies^8,10^. We calculated the enrichment of GenAge genes for each subnetwork (see Table S5 for the results). The most significant overlap was seen in the hotpink^old^ (p-value = 1.15E-2). This subnetwork is relatively small (107 genes) and is highly enriched for cell cycle genes (adjusted p-value = 1.3E-52). The second most significant subnetwork was the turquoise^old^, which shared 34 common genes with GenAge (p-value=1.26E-2) (see Table S5 and Table S6). It is of note that turquoise^old^ contains *APOE* (apolipoprotein E), which is one of the very few genes that are reproducible in genome-wide association studies for human longevity^26^.

*Criterion 2. Enrichment for differential expressed aging genes*: We applied DESeq^27^ to identify differentially expressed genes (DEGs) between young and old adipose tissue samples. At FDR 0.05, there were 382 DEGs, among which 265 were protein-coding genes with 148 up-regulated and 117 down-regulated (Table S7). We also performed sample down-sampling to ensure that the DEGs were robust and did not dependent on the difference of sample size (see Supplementary Results). The up-regulated protein coding DEGs are significant enriched for functions like “Extracellular matrix” (FDR 6.74E-6), “glycoprotein” (FDR 1.05E-5), and “signal peptide” (FDR 2.43E-3), and the down-regulated genes are significant enriched for Transmembrane (FDR 2.56E-3) and so on (Table S8). It is of note that the top ranked DEG was *CDKN2A*, whose expression levels increased more than threefold from young to old adipose tissues (FDR 1.75E-6). *CDKN2A* encodes multiple isoforms and the isoform, p16^INK4A^, is a commonly used biomarker for cellular senescence^28^. Studies have shown that by clearing p16^INK4A^-positive cells, mice could have a longer lifespan with delayed onset of age-related disorders^29^.

We mapped the aging DEGs to the old adipose tissue co-expression network and summarized the results in Table 1. There are 3 modules significantly enriched for the aging DEGs, i.e., turquoise^old^ (100 common genes with p-value 3.55E-44), magenta^old^ (31 genes with p-value 3.51E-15), and yellow^old^ (28 genes with p-value 4.18E-6). Turquoise^old^ contained about two fifth of all the significant aging DEGs (100 out of 265), indicating aging had the strongest influence on gene expression to this subnetwork compared to other subnetworks.

**Table 1.**
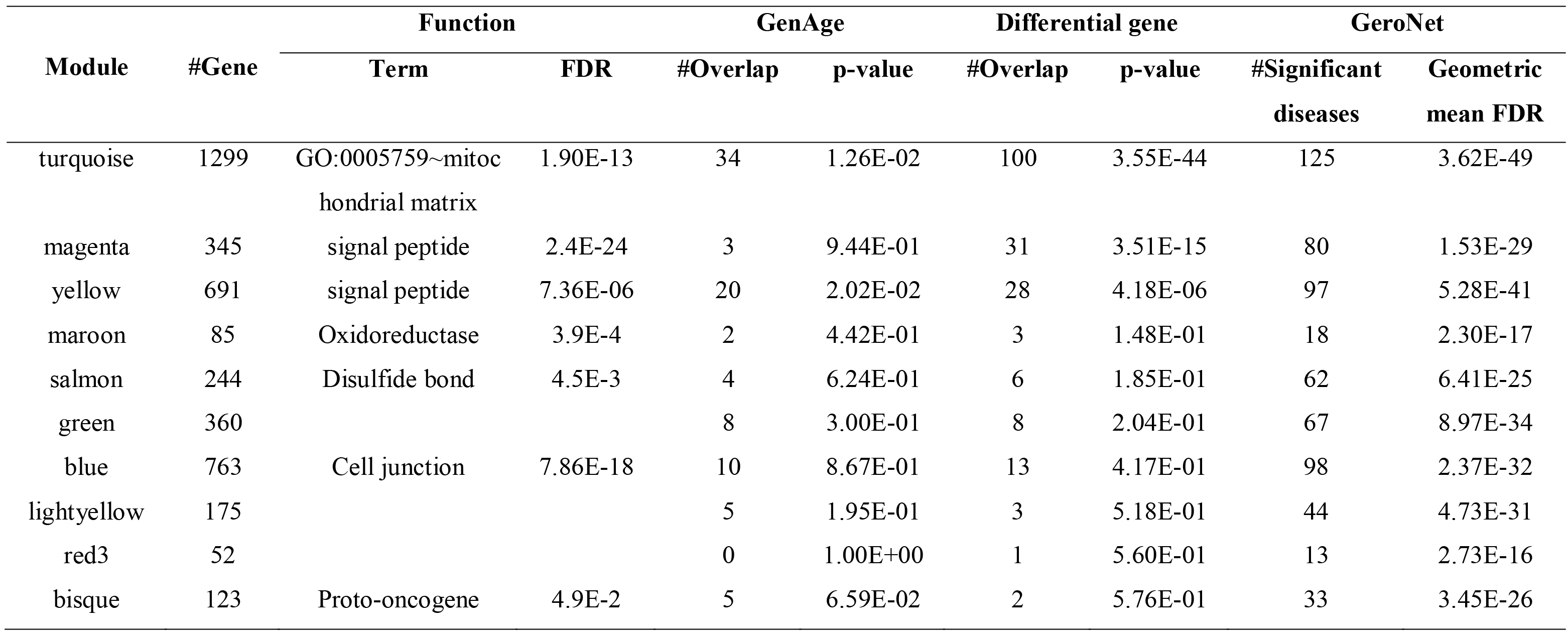
A summary of the top ten largest modules in old adipose tissue and their enrichment for aging DEGs, GenAge genes, and the significance for connecting to ARDs as indicated by GeroNet

*Criterion 3. Enrichment for disease genes*: We previously manually curated genes associated with 277 diseases and traits, among which 84 diseases and traits had at least 20 associated genes^12^. We evaluated the enrichment of disease genes in each subnetwork for all the 84 diseases/traits. As shown in Fig 3, there exists significant enrichment of disease genes in multiple subnetworks. For example, the greenyellow^old^ subnetwork overlapped with multiple immune related diseases (e.g., vitiligo, ulcerative colitis, inflammatory bowel disease, and Alzheimer’s disease) while it is highly enriched for “immunity” (adjusted p-value = 4.3E-34). The tuquoise^old^ subnetwork significantly overlapped with genes associated with obesity-related traits, Parkinson’s disease, metabolite traits, and was the only subnetwork that significantly overlapped with type 2 diabetes genes, indicating it may play an important function in mediating the development of these metabolic related diseases.

**Fig 3.**
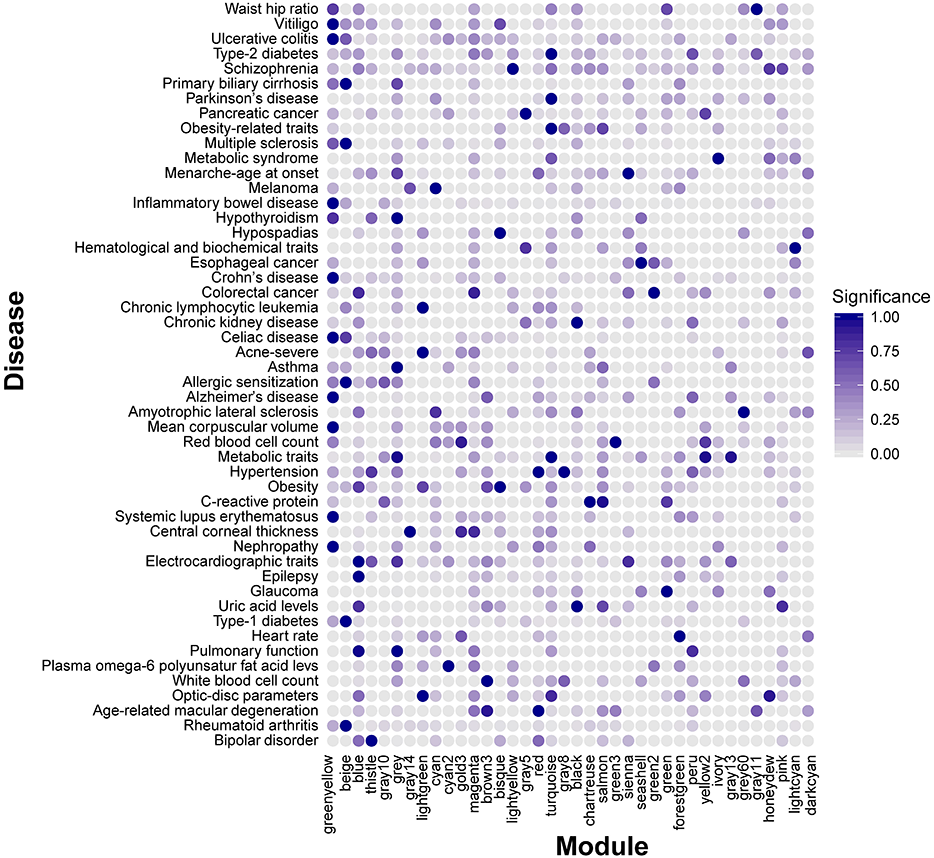
Enrichment of disease genes for each subnetwork derived from old adipose tissues. Only subnetworks/diseases having at least one significant enrichment (p-value <0.02) are shown.

*Criterion 4. Strong connection with age-related diseases by GeroNet calculation*: GeroNet is a method that allows the consideration of connections between aging and disease genes through network connections. Different from criteria 3, it considers indirect interactions (i.e., interactions mediated by other genes) between aging and disease genes^12^. We performed GeroNet analysis using GenAge genes as aging genes and disease genes from 277 disease and trait categories. We used the HPRD (human protein reference database) as a reference PPI network^12^. The results from GeroNet showed that turquoise^old^ was the most important subnetwork in mediating the connections between aging and multiple diseases in all the 85 subnetworks. This result further suggests that turquoise^old^ subnetwork plays a critical role in mediating the connections between aging and age-related disease including metabolic diseases such as type 2 diabetes.

*Criterion 5. Enrichment for differential co-expressions*: As we have shown in the previous section, we observed profound co-expression re-wiring between magenta^young^ and turquoise^old^ modules. To assess the significance level of re-wiring, we performed a permutation analysis. Briefly, for a subnetwork of size *n*, we randomly selected *n* genes from all the genes in the transcriptome to form a random subnetwork. Using the same set of young and old tissue samples, we calculated the number of significant differential co-expressions for this “random” subnetwork between young and old ages. By running this permutation test for 100 times, we estimated a p-value for observing the actual number of differentially co-expressed gene-gene pairs in our real data. Using this permutation estimation, the estimated p-value for observing 1,052 differential co-expressions was close to 0 (p <= 1E-324), which indicates that co-expression rewiring in the turquoise^old^ module was highly significant.

To summarize, we listed the results for the top 10 modules with more than 50 protein coding genes in Table 1 (sorted based on the enrichment for differentially expressed aging genes). The turquoise^old^ module was ranked at the 1^st^ position for enrichment of aging DEGs, it was also ranked at 2^nd^ for enrichment of GenAge genes and 1^st^ based on GeroNet computation. Collectively, the turquoise^old^ stood out as a significant subnetwork in which aging and a group of metabolic diseases like type 2 diabetes are connected.

To visualize the turquoise^old^ subnetwork, we mapped its genes onto the HPRD protein-protein interaction (PPI) network and displayed the largest connected component in Fig 4A. We annotated the diabetes genes (square shaped nodes), significantly DEGs between young and old samples (nodes in red color for up-regulated and green color for down-regulated gene expressions in old adipose tissue samples) and *APOE*-perturbed genes (nodes with thick border, the *APOE* knockout DEGs were derived from GSE44653) in Fig 4a. It is of note that signaling pathways important in growth regulation and metabolic function such as the *PPARG* and *GHR/AKT* pathway are contained in this PPI subnetwork. Similarly, we show the greenyellow^old^ PPI network in Fig 4B and genes associated with inflammatory diseases are highlighted with thicker borders.

**Fig 4.**
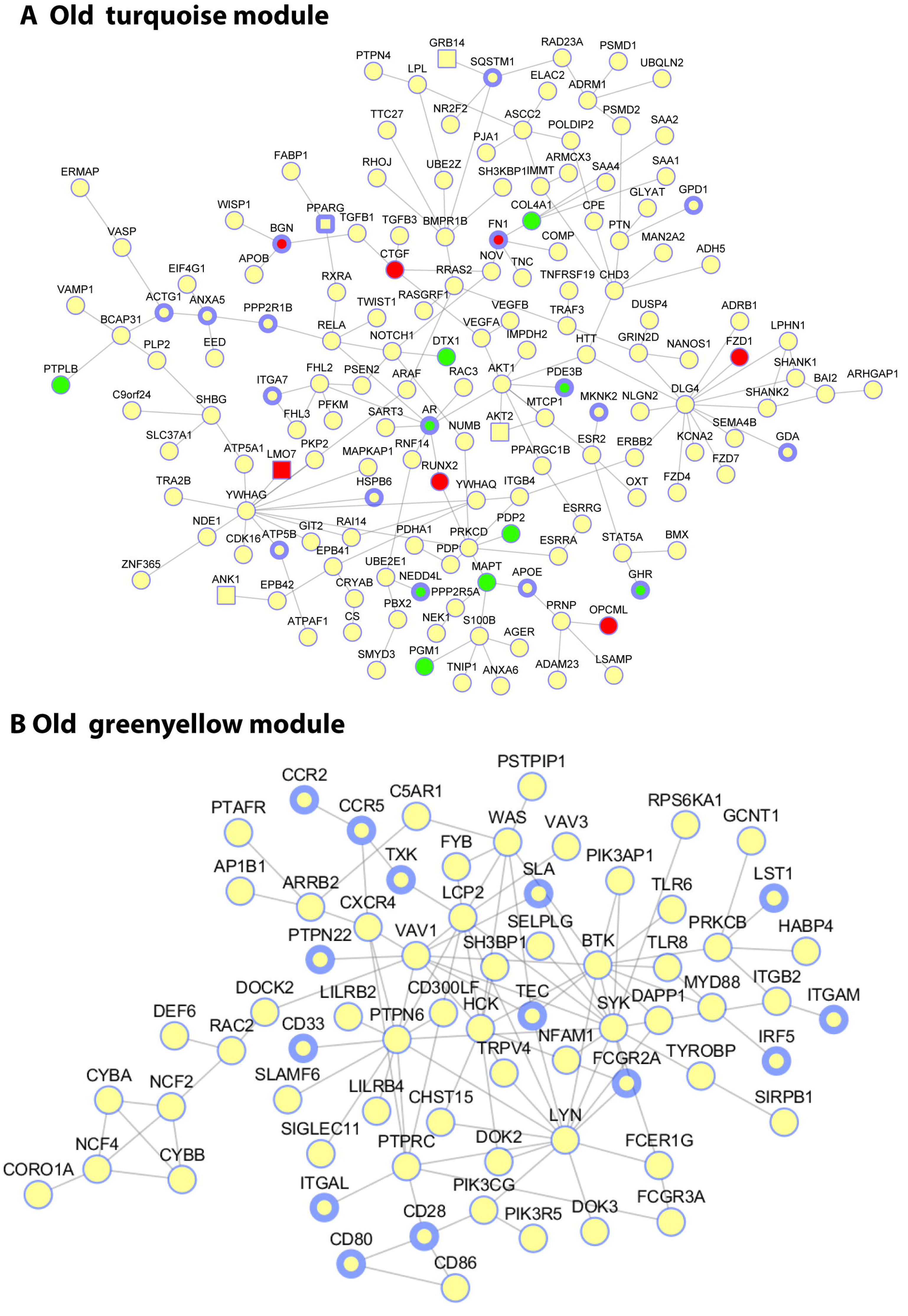
A protein-protein interaction network view of two subnetworks derived from old adipose tissue. (A) turquoise^old^ module and its association to aging, diabetes, and *APOE* perturbed genes. Each node denotes a gene and each edge denotes a protein-protein interaction. The nodes in square shape (e.g., *AKT2*) are diabetes associated genes obtained from GWAS catalog and OMIM; the nodes with filled red-(up) or green-(down) (e.g., *MAPT*) are significantly differentially expressed genes (at FDR <= 0.05 by DESeq) between young and old samples; the nodes with blue thick border (e.g., *APOE* and *GDA*) are *APOE* perturbed genes as defined in CREEDS. (B) greenyellow^old^ module mapped to PPI network and genes associated with inflammatory diseases (nodes with thicker border), including vitiligo, ulcerative colitis, multiple sclerosis, inflammatory bowel disease, Crohn’s disease, celiac disease, Alzheimer’s disease, and systemic lupus erythematosus.

### Prioritize drugs and gene targets using a network pharmacogenomic approach

An important goal of human aging and geroscience research is to discover effective interventions to slow aging and delay the onset of age-related diseases. As our analysis first prioritizes aging subnetworks that are closely related to diseases, the next step is to identify drugs or target genes that may reverse the changes in the prioritized subnetworks. To achieve this goal, we adopt a network pharmacogenomic approach based on the concept of connectivity maps (CMap)^15^. The goal of CMap is to identify drug or gene perturbations that cause gene expression changes in the opposite direction as seen in the disease condition. The identified drugs or genes are considered to have a good chance of treating the disease. The CMap concept has received several proof-of-principle validations and an increasing number of successful applications have been reported^17,30^. These successes stimulated multiple large efforts to construct large scale perturbation expression signature databases, e.g., The Library of Network-Based Cellular Signatures (LINCS) Program^31^, and a crowd extracted expression of differential signatures (CREEDS)^32^. Such databases are valuable resources to combine with the aging network biology to make new geroprotector discoveries.

For our study, we hypothesize that if a drug or gene perturbation could reverse aging DEGs in our prioritized aging subnetworks, then this drug/gene is likely to be geroprotective. To implement this approach, we divided the 265 differential genes into 148 up-regulated genes and 117 down-regulated genes (Table S7) and use them to query the drug/gene perturbation databases. The up-regulated aging genes were mostly enriched for “extracellular matrix” with FDR 6.7E-6 and the down-regulated aging genes were mostly enriched for “transmembrane” with FDR 1.48E-3.

We relied on CREEDS database which is a manually curated list of 4,295 single-drug perturbations and 8,620 single-gene perturbations generated by leveraging the public available gene expression data deposited in the Gene Expression Omnibus^32^. Specifically, we screened the CREEDS to identify either drug or gene perturbations that could reverse the aging DEGs (i.e. perturbations that up-regulate those repressed aging gene expressions or down-regulate those elevated aging gene expressions). We performed a two-way comparison between drug/gene perturbation signatures with the up-and down-regulation of the aging gene expression signatures separately. We only show the results for analyzing the down-regulated aging signatures. We leave the results for analyzing the up-regulated aging signatures in the supplementary material (Dataset 2). We ranked drug and gene perturbations based on the p-values of the Fisher’s exact test for overlap between the perturbation signatures and the aging signatures. We considered the aging DEGs restricted to the turquoise^old^ module, given this subnetwork is of particular interest as a converging point of aging and diseases.

*Top drugs based on differential aging genes*: We list the top 5 identified drugs (among 4,295) in Table 2. It is of note that same drugs may appear more than once. This happens when the top signatures from different experiments tested the same drugs, or when the top signatures were from the same drug experiment but based on comparing different samples, which were treated as unique signatures by CREEDS. We include metformin as a reference due to its therapeutic applications in multiple disease areas. Due to its robust record of safety, metformin is among the top candidate human geroprotectors that have been cleared by FDA to enter clinical trials^33^.

**Table 2.**
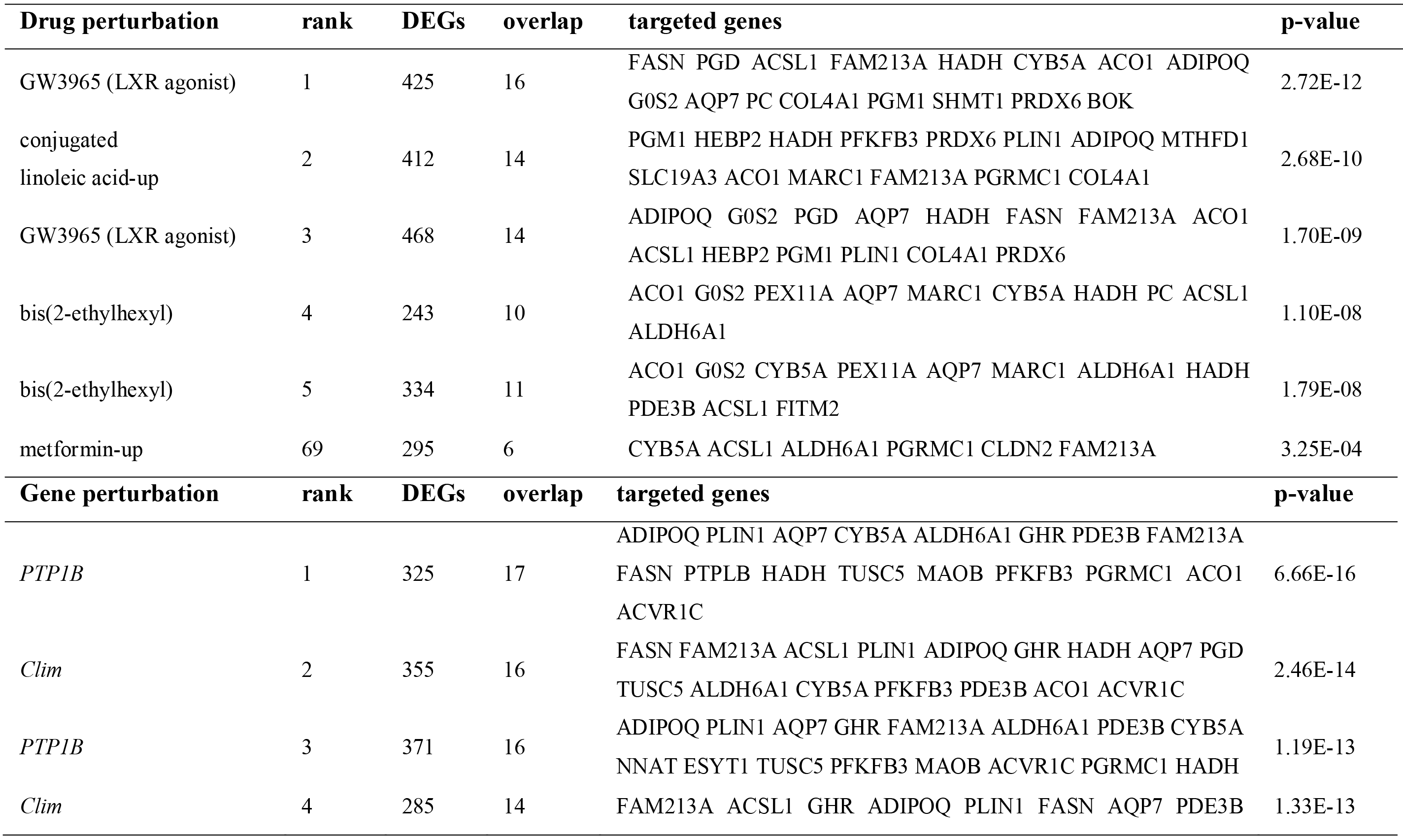
Top drug and gene perturbations that up-regulate the down-regulated aging DEGs in the turquoise^old^ module for adipose tissue.

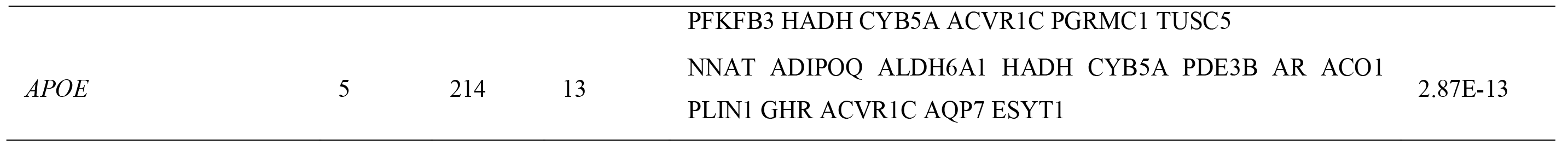

As Table 2 shows, several conjugated linoleic acid (CLA) induced gene expression signatures were ranked at the top. These signatures were derived from a study investigating the effect of CLA supplementation on gene expression in human adipose tissue biopsies (GSE16615)^34^. From the enrichment analysis, both isomers of CLA, either cis-9, trans-11 CLA or trans-10, cis-12 CLA, or their mixture up-regulated the expression of multiple genes that were down-regulated in the old GTEx adipose tissue. These genes encompass functions such as enzymes in various metabolic pathways (*PFKFB3*, *PGM1*, *GLUL*, *MTHFD1*, *ACO1* and *HADH*), cofactor and metabolite transporters (*SLC19A3* and *AQP7*), redox homeostasis (*FAM213A* and *PRDX6*), *PPAR* signaling pathway genes (*ADIPOQ*), cell cycle regulation (*CDKN2C*), and lipid metabolism (*PLIN1* and *HADH*). The function of these genes is generally consistent with the reported favorable effects of CLA on cancer, atherosclerosis, body weight and fat mass in animal models.

Another distinct hit is the liver X receptor (LXR) agonist GW3965 and the perturbation signature was based on a study to investigate LXR’s role on regulating glucose uptake in human adipocytes (GSE41223)^35^. The canonical role of LXR is regulation of reverse cholesterol transport and cholesterol efflux, but its role also extends to the regulation of glucose and fatty acid metabolism and a wide variety of endocrine processes^36^. The targeted genes that overlap with the aging DEGs are consistent with their function in lipid synthesis (*FASN* and *ACSL1*), glucose metabolism (*PGD*, *PC* and *PGM1*), as well as other metabolic pathways (*CYB5A*, *HADH*, *ACO1*, *AQP*7 and *SHMT1*).

In addition, metformin ranked at 65^th^ (69^th^ by turquoise^old^ module) out of a total of 4,295 drugs (in top 2%). This drug has been shown to be able to increase both lifespan and health span in mice^37^.

*Top single gene perturbations that strongly enriched for differential aging gene expression*: We summarized the top 5 identified gene perturbations (among 8,620) whose up-regulated genes significantly overlapped with the 117 down-regulated aging DEGs in Table 2.

The top gene perturbation signature is from a tyrosine-protein phosphatase non-receptor type 1 (PTP1B, encoded by *PTPN1* gene in human) knockout mouse mammary gland tissues^38^. *PTP1B* is a negative regulator of both insulin and leptin signaling and is involved in the control of glucose homeostasis and energy expenditure^39^. *PTP1B* deficient mice show increased insulin sensitivity and resistance to diet-induced obesity^40^. *PTPN1* is a therapeutic target for type 2 diabetes, obesity and cancer. The second of our top gene perturbations is the Co-factor of the LIM domains (Clim), that was studied using a dominant negative Clim that interfered with function of both Clim1 and Clim 2 (Ldb2 and Ldb1 respectively)^41^. Interestingly, Clim2 is known to interact with *LMO4*, which is a metabolic responsive inhibitor of *PTP1B* that controls hypothalamic leptin signaling. More interestingly, study showed that *LXR* agonists could down-regulate *PTP1B* gene expression and ameliorate TNF alpha-induced insulin resistance in murine brown adipocytes^42^. Considering all the connections among the top hits, a signaling map among the top genes emerges and is shown in Fig S3. The signaling pathway inferred by ANDRU further suggests that the gene hits we found are unlikely random false positives, given the meaningful connections among them.

The *APOE* knockout DEG signature derived from mouse fat (GSE44653) was ranked at 5^th^. *APOE* is one of the two reproducible longevity genes for human, and is also known to be associated to many diseases like Alzheimer’s disease. APOE is a multi-functional protein expressed highly in adipose tissue and plays significant role in modulating lipid metabolism. The highlighted drugs and genes appear to target on an overlapping gene set that includes *ADIPOQ*, the gene encoding adiponectin. Adiponectin is an adipokine or hormone that originates from adipose tissue. Although debated, adiponectin is generally considered to have beneficial effects on ARDs such as obesity and type 2 diabetes.

### The prioritized drugs and genes show tissue specificity when ANDRU is applied to another tissue

It is known that aging and ARDs show tissue specificity^6,7^. To evaluate if the ranking of drugs or genes would depend on tissue types, we applied ANDRU to a second tissue type, namely, the artery aorta. The results are summarized in Dataset 3. There were 34 young samples (with mean age and standard deviation of 27±4.5) and 33 old samples (with mean age and standard deviation of 67±1.1) used by ANDRU. At FDR of 0.05, 459 up-regulated and 325 down-regulated differential protein-coding genes were identified by comparing the young vs. old samples. These DEGs are significantly enriched for glycoprotein and several other categories. For the drug and gene perturbation prediction, the top 3 drug perturbations that up-regulate the down-regulated aging DEGs are N-methyl-D-aspartate (NMDA) antagonists. The NMDA receptor has been well-studied for its association with brain aging^43,44^. Next to the NMDA antagonists, curcumin was ranked at 4,5,6,8, and 10 in the list. Curcumin is a chemical produced naturally by some plants like ginger root. Curcumin and its metabolite, tetrahydrocurcumin (THC), has been experimentally validated to increase lifespans of nematode roundworm, fruit fly, and mouse^45^. It is also known that curcumin has anti-oxidative, anti-lipofusinogenesic, and anti-aging effects in the brain^46^ and might be effective in treating diseases caused by low grade inflammation including cancer and Alzheimer’s disease^47^. From network analysis, curcumin was also ranked at the top for aging DEGs in the yellow subnetwork (919 protein coding genes enriched for Zinc finger regions and DNA binding) in old artery aorta tissue (Dataset 3). The top gene perturbations that predicted to be associated with the aging in artery aorta including *TGFBR1*, *RB1*, *TP53*, *STAT1*, etc., which are well-known to be involved in aging process^25^. It is of note that the prioritized drugs are different for the two tissue types we evaluated, implying the underlying aging-related changes and how they interconnect with diseases are tissue specific.

## Discussions

In this work, we demonstrated a novel informatics pipeline ‑ ANDRU to construct human aging network and identify interventions that can be geroprotective. We applied ANDRU to GTEx adipose and artery tissues. With aging, adipose tissue undergoes significant changes and contributes to the development of insulin resistance, metabolic dysfunction, inflammation, and impaired regenerative capacity^22^. From more than eighty adipose subnetworks, we prioritized and highlighted the turquoise^old^ for its putative role as the interface between aging and several ARDs. Since different subnetworks often have distinct functions and show differential connections to various diseases, our approach provides the option to reverse the aging in different subnetworks to address different types of diseases (e.g., the inflammation related diseases vs. metabolic related diseases). Since the diseases of concern often vary among individuals, the option of providing prioritization based on aging subnetworks considering individual’s genetic and genomic profiles could facilitate personalized geroprotection in future.

The importance and feasibility of performing *in silico* drug screening for geroprotection has been first demonstrated by Zhavoronkov et al. followed by some experimental validations^48^. Our method is conceptual similarity to their method but has a very different way of implementation. For example, we relied on a *de novo* network construction approach to build aging networks while GeroScope used by Zhavoronkov et al. considered prior knowledge of known aging pathways. While both approaches considered age-related transcriptional changes, the input dataset and methods used to incorporate such information were different. Therefore, our approach represents a significant alternative option for performing *in silico* geroprotector discovery.

The current implementation of ANDRU has its limitations and will require future improvement. First, as we relied on existing gene expression signature databases, we are limited to the perturbations collected by these databases and important genes or drugs could be missed if the data was not generated or not included into the database. Although this “incompleteness” of perturbation signatures will likely persist in the near future, perturbation datasets are continuing to grow. For example, the NIH LINCS project is generating high throughput perturbation expression signature data at very large scale^31^, making it another useful resource for the network pharmacogenomics development. Second, CREEDS contains data collected from multiple species, and the non-human data or data obtained from different tissue types could cause either false positive or false negative hits. This would require additional conformational experiments to ensure perturbation signatures are reproducible in the desirable tissue types.

Finally, in this pilot work, we considered only two tissue types to illustrate the pipeline. From these two tissues, ANDRU output different drug/gene candidates. This suggests that to achieve optimized systemic geroprotection, combining several drugs should be something for consideration. As GTEx profiles more than 50 tissue types, it is a future work to investigate aging networks and search for geroprotectors at a “whole-body” scale. With an unprecedented increasing amount of human omics data, we believe that integrating them to refine the aging networks, and leveraging them to search for the most effective human geroprotectors represents a promising strategy to speed up the geroscience research.

## Experimental Procedures

### Data collection and processing

Human gene expression data (both fpkm and raw reads count) is from GTEx portal^20^. We divided individuals into three groups according to their chronological ages: young group (age <=35), old group (age >=65), and those in between. We only considered young and old groups for further analysis. We performed a few data processing steps to require genes to have at least 0.1 RPKM in 2 or more individuals followed by a quantile normalization across genes. Following GTEx study^20^, we adjusted the following confounding factors: (1) gender, (2) collection center, (3) RIN, (4) ischemic time, and (5) 3 genotype principal components. The genotype PCs were constructed using GCTA^49^ with several quality control steps using plink (e.g., -- MAF 0.1, -- geno 0.05, --hwe 1e-6, --Chr 1-22).

### Construction of young and old gene co-expression networks

For both young and old samples, the expression values for each gene were quantile normalized into a standard normal distribution. We then removed sample outliers by a hierarchical clustering of the samples based on the expression data for young and old groups respectively. 36 samples in young and 52 samples in old group for adipose tissue and 34 samples in young and 33 samples in old for artery aorta were kept to construct young and old networks by weighted gene correlation network analysis (WGCNA)^21^. We set the soft power to be 5 for adipose and 4 for artery aorta, respectively.

### Differential gene expression and differential connectivity analysis

We collected the read counts for young and old samples separately and applied two pipelines HTSeq/DESeq and HTSeq/edgeR to call the differential genes between young and old samples using adjusted p-value less than 0.05 as threshold. We applied DGCA to perform differential connectivity analysis^23^. DGCA computes and analyses differential correlations between gene pairs across young and old samples. A gene pair was called differentially connected if p-value calculated by DGCA was less than or equal to 5E-4. We then performed a permutation analysis to evaluate the enrichment of differential connected gene pairs in each module.

### Drug and perturbed gene signatures

We considered the signatures of 4,295 drugs and 8,620 gene perturbations collected in CREEDS for this study^32^. We performed enrichment analysis between perturbation signatures with aging DEGs by Fisher’s exact test. A drug was highly ranked if drug induced down-regulated genes significantly overlapped with up-regulated aging DEGs, or drug induced up-regulated genes significantly overlapped with down-regulated aging DEGs. Gene perturbations were similarly ranked.

## Acknowledgments

JY is currently supported by a postdoctoral fellowship from Unity Biotechnology. ZT receives financial support from Unity Biotechnology as a consultant. This work is partially supported by NIH/NIA R01-AG055501 and a Leducq foundation awarded to ZT. This work was also supported in part through the computational resources and staff expertise provided by Scientific Computing at the Icahn School of Medicine at Mount Sinai.

### Contributions

ZT conceived the concept of the work. JY and ZT performed the analysis. JY, SH, YS, ZT wrote the manuscript. All authors helped to discuss and improve the work.

### Competing Interests

The authors have declared no competing interests.

## Supporting Information

**Supplementary Results**

**Supplementary Figures**

**Supplementary Tables**

**Supplementary Datasets**

## References

1 Kennedy, B. K. et al. Geroscience: linking aging to chronic disease. Cell 159, 709–713, doi:10.1016/j.cell.2014.10.039 (2014).

2 Everitt, A. V. et al. Dietary approaches that delay age-related diseases. Clinical interventions in aging 1, 11–31 (2006).

3 Moskalev, A. et al. Geroprotectors.org: a new, structured and curated database of current therapeutic interventions in aging and age-related disease. Aging (Albany NY) 7, 616, doi:10.18632/aging.100799 (2015).

4 Barardo, D. et al. The DrugAge database of aging-related drugs. Aging cell, doi:10.1111/acel.12585 (2017).

5 Kumar, S. & Lombard, D. B. Finding Ponce de Leon’s Pill: Challenges in Screening for Anti-Aging Molecules. F1000Research 5, 406, doi:10.12688/f1000research.7821.1 (2016).

6 Glass, D. et al. Gene expression changes with age in skin, adipose tissue, blood and brain. Genome Biology 14, 1–12, doi:10.1186/gb-2013-14-7-r75 (2013).

7 Yang, J. et al. Synchronized age-related gene expression changes across multiple tissues in human and the link to complex diseases. Scientific reports 5, 15145, doi:10.1038/srep15145 (2015).

8 Zhang, Q. et al. Systems-level analysis of human aging genes shed new light on mechanisms of aging. Human Molecular Genetics, doi:10.1093/hmg/ddw145 (2016).

9 Fernandes, M. et al. Systematic analysis of the gerontome reveals links between aging and age-related diseases. Human Molecular Genetics, doi:10.1093/hmg/ddw307.

10 Wang, J., Zhang, S., Wang, Y., Chen, L. & Zhang, X. S. Disease-aging network reveals significant roles of aging genes in connecting genetic diseases. PLoS computational biology 5, e1000521, doi:10.1371/journal.pcbi.1000521 (2009).

11 Johnson, S. C., Dong, X., Vijg, J. & Suh, Y. Genetic evidence for common pathways in human age-related diseases. Aging cell 14, 809–817, doi:10.1111/acel.12362 (2015).

12 Yang, J. et al. Discover the network mechanisms underlying the connections between aging and age-related diseases. 6, 32566, doi:10.1038/srep32566 (2016).

13 Johnson, S. C. et al. Network analysis of mitonuclear GWAS reveals functional networks and tissue expression profiles of disease-associated genes. Human Genetics 136, 55–65, doi:10.1007/s00439-016-1736-9 (2016).

14 Csermely, P., Korcsmáros, T., Kiss, H., London, G. & Nussinov, R. Structure and dynamics of molecular networks: A novel paradigm of drug discovery A comprehensive review. Pharmacology & Therapeutics 138, 333–408, doi:10.1016/j.pharmthera.2013.01.016 (2013).

15 Lamb, J. Innovation ‑ The Connectivity Map: a new tool for biomedical research. Nat Rev Cancer 7, 54–60 (2007).

16 Iorio, F. et al. Discovery of drug mode of action and drug repositioning from transcriptional responses. Proceedings of the National Academy of Sciences 107, 14621–14626, doi:10.1073/pnas.1000138107 (2010).

17 Dudley, J. T. et al. Computational Repositioning of the Anticonvulsant Topiramate for Inflammatory Bowel Disease. Science Translational Medicine 3, doi:10.1126/scitranslmed.3002648 (2011).

18 Wagner, A. et al. Drugs that reverse disease transcriptomic signatures are more effective in a mouse model of dyslipidemia. Molecular Systems Biology 11, 791, doi:10.15252/msb.20145486 (2015).

19 Calvert, S. et al. A network pharmacology approach reveals new candidate caloric restriction mimetics in C. elegans. Aging Cell 15, 256–266, doi:10.1111/acel.12432 (2016).

20 The GTEx Consortium. The Genotype-Tissue Expression (GTEx) pilot analysis: Multitissue gene regulation in humans. Science 348, 648–660, doi:10.1126/science.1262110 (2015).

21 Zhang, B. & Horvath, S. A general framework for weighted gene co-expression network analysis. Stat Appl Genet Mo B 4 (2005).

22 Palmer, A. K. & Kirkland, J. L. Aging and adipose tissue: potential interventions for diabetes and regenerative medicine. Experimental gerontology 86, 97–105, doi:10.1016/j.exger.2016.02.013 (2016).

23 McKenzie, A. T., Katsyv, I., Song, W. M., Wang, M. H. & Zhang, B. DGCA: A comprehensive R package for Differential Gene Correlation Analysis. Bmc Syst Biol 10 (2016).

24 Tchkonia, T. et al. Fat tissue, aging, and cellular senescence. Aging cell 9, 667–684, doi:10.1111/j.1474-9726.2010.00608.x (2010).

25 Tacutu, R. et al. Human Ageing Genomic Resources: Integrated databases and tools for the biology and genetics of ageing. Nucleic Acids Res 41, D1027–D1033, doi:Doi 10.1093/Nar/Gks1155 (2013).

26 Schachter, F. et al. Genetic Associations with Human Longevity at the Apoe and Ace Loci. Nat Genet 6, 29–32, doi:Doi 10.1038/Ng0194-29 (1994).

27 Anders, S. & Huber, W. Differential expression analysis for sequence count data. Genome Biology 11, doi:Artn R10610.1186/Gb-2010-11-10-R106 (2010).

28 Sharpless, N. E. & Sherr, C. J. Forging a signature of in vivo senescence. Nature reviews. Cancer 15, 397–408, doi:10.1038/nrc3960 (2015).

29 Baker, D. J. et al. Naturally occurring p16(Ink4a)-positive cells shorten healthy lifespan. Nature 530, 184–+ (2016).

30 Mirza, N., Sills, G. J., Pirmohamed, M. & Marson, A. G. Identifying New Antiepileptic Drugs Through Genomics-Based Drug Repurposing. Human molecular genetics 26, doi:10.1093/hmg/ddw410 (2017).

31 Duan, Q. et al. L1000CDS2: LINCS L1000 characteristic direction signatures search engine. npj Systems Biology and Applications 2, 16015, doi:10.1038/npjsba.2016.15 (2016).

32 Wang, Z. C. et al. Extraction and analysis of signatures from the Gene Expression Omnibus by the crowd. Nat Commun 7 (2016).

33 Barzilai, N., Crandall, J. P., Kritchevsky, S. B. & Espeland, M. A. Metformin as a Tool to Target Aging. Cell Metabolism 23, 1060–1065, doi:10.1016/j.cmet.2016.05.011 (2016).

34 Herrmann, J. et al. Isomer-specific effects of CLA on gene expression in human adipose tissue depending on PPARgamma2 P12A polymorphism: a double blind, randomized, controlled cross-over study. Lipids in health and disease 8, 35, doi:10.1186/1476-511X-8-35 (2009).

35 Pettersson, A. M. L. et al. LXR is a negative regulator of glucose uptake in human adipocytes. Diabetologia 56, 2044–2054, doi:10.1007/s00125-013-2954-5 (2013).

36 Calkin, A. C. & Tontonoz, P. Transcriptional integration of metabolism by the nuclear sterol-activated receptors LXR and FXR. Nature reviews. Molecular cell biology 13, 213–224, doi:10.1038/nrm3312 (2012).

37 Martin-Montalvo, A. et al. Metformin improves healthspan and lifespan in mice. Nat Commun 4 (2013).

38 Milani, E. S. et al. Protein tyrosine phosphatase 1B restrains mammary alveologenesis and secretory differentiation. Development (Cambridge, England) 140, 117–125, doi:10.1242/dev.082941 (2013).

39 Tsou, R. C. & Bence, K. K. The genetics of PTPN1 and obesity: insights from mouse models of tissue-specific PTP1B deficiency. Journal of obesity (2012).

40 Elchebly, M., Payette, P. & Michaliszyn, E. Increased insulin sensitivity and obesity resistance in mice lacking the protein tyrosine phosphatase-1B gene. …, doi:10.1126/science.283.5407.1544 (1999).

41 Salmans, M. L. et al. The Co-factor of LIM Domains (CLIM/LDB/NLI) Maintains Basal Mammary Epithelial Stem Cells and Promotes Breast Tumorigenesis. PLoS Genetics 10, doi:10.1371/journal.pgen.1004520 (2014).

42 Fernández-Veledo, S., Nieto-Vazquez, I., Rondinone, C. M. & Lorenzo, M. Liver X receptor agonists ameliorate TNFalpha-induced insulin resistance in murine brown adipocytes by downregulating protein tyrosine phosphatase-1B gene expression. Diabetologia 49, 3038–3048, doi:10.1007/s00125-006-0472-4 (2006).

43 Magnusson, K. R., Brim, B. L. & Das, S. R. Selective Vulnerabilities of N-methyl-D-aspartate (NMDA) Receptors During Brain Aging. Frontiers in aging neuroscience 2, 11, doi:10.3389/fnagi.2010.00011 (2010).

44 Foster, T., Kyritsopoulos, C. & Kumar, A. Central role for NMDA receptors in redox mediated impairment of synaptic function during aging and Alzheimer’s disease. Behav Brain Res 322, 223–232, doi:10.1016/j.bbr.2016.05.012 (2017).

45 Shen, L. R., Parnell, L. D., Ordovas, J. M. & Lai, C. Q. Curcumin and aging. Biofactors 39, 133–140, doi:10.1002/biof.1086 (2013).

46 Bala, K., Tripathy, B. C. & Sharma, D. Neuroprotective and anti-ageing effects of curcumin in aged rat brain regions. Biogerontology 7, 81–89, doi:10.1007/s10522-006-6495-x (2006).

47 Sikora, E., Bielak-Zmijewska, A., Mosieniak, G. & Piwocka, K. The Promise of Slow Down Ageing May Come from Curcumin. Curr Pharm Design 16, 884–892, doi:Doi 10.2174/138161210790883507 (2010).

48 Aliper, A. et al. In search for geroprotectors: in silico screening and in vitro validation of signalome-level mimetics of young healthy state. Aging 8, 2127–2152, doi:10.18632/aging.101047 (2016).

49 Yang, M. et al. Microvesicles secreted by macrophages shuttle invasion-potentiating microRNAs into breast cancer cells. Molecular cancer 10, 117, doi:10.1186/1476-4598-10-117 (2011).

